# Genetic Analysis of Substrain Divergence in NOD Mice

**DOI:** 10.1101/013037

**Authors:** Petr Simecek, Gary A. Churchill, Hyuna Yang, Lucy B. Rowe, Lieselotte Herberg, David V. Serreze, Edward H. Leiter

## Abstract

The NOD mouse is a polygenic model for type 1 diabetes that is characterized by insulitis, a leukocytic infiltration of the pancreatic islets. During ~35 years since the original inbred strain was developed in Japan, NOD substrains have been established at different laboratories around the world. Although environmental differences among NOD colonies capable of impacting diabetes incidence have been recognized, differences arising from genetic divergence have not previously been analyzed. We use both Mouse Diversity Array and Whole Exome Capture Sequencing platforms to identify genetic differences distinguishing 5 NOD substrains. We describe 64 SNPs, and 2 short indels that differ in coding regions of the 5 NOD substrains. A 100 kb deletion on Chromosome 3 distinguishes NOD/ShiLtJ and NOD/ShiLtDvs from 3 other substrains, while a 111 kb deletion in the *Icam2* gene on Chromosome 11 is unique to the NOD/ShiLtDvs genome. The extent of genetic divergence for NOD substrains is compared to similar studies for C57BL6 and BALB/c substrains. As mutations are fixed to homozygosity by continued inbreeding, significant differences in substrain phenotypes are to be expected. These results emphasize the importance of using embryo freezing methods to minimize genetic drift within substrains and <u>of applying appropriate genetic nomenclature to permit substrain recognition when one is used.</u>

## INTRODUCTION

The NOD mouse represents a premier animal model for the study of spontaneous insulitis and autoimmune Type 1 diabetes. This inbred strain, first reported in 1980, was developed in Japan by selective breeding of outbred ICR:Jcl mice at the Shionogi Research Laboratories (Makino et al. 1980). The Central Laboratory for Experimental Animals (CLEA Japan) began receiving breeding stock from the NOD/Shi source colony for international distribution by 1986. However, prior to that time, NOD/Shi breeding stock from various sources had been obtained in two locations in the United States, one in Germany, and one in Australia (reviewed in (Leiter 1998)). Accumulation of new mutations fixed to homozygosity by inbreeding can be expected to produce significant substrain divergence over time, and potentially, differences in substrain characteristics. When NOD mice are maintained by inbreeding for at least 10 generations separately from the source colony (NOD/Shi), they are designated as substrains and receive either the colony holder’s and/or institution’s symbol. Among the currently most studied NOD substrains are NOD/ShiJcl (CLEA Japan, Inc., http://www.clea-japan.com/en/animals/animal_b/b_06.html), NOD/ShiLtJ (The Jackson Laboratory, http://jaxmice.jax.org/strain/001976.html) and the NOD/ShiLtDvs substrain derived from it, NOD/MrkTac (Taconic, http://www.taconic.com/wmspage.cfm?parm1=871), and NOD/BomTac.

Phenotypic differences between and within NOD substrains have been observed (De Riva et al. 2013; Leiter 1993; Takayama et al. 1993). Given the strong role that environmental factors <u>and especially the microbiome</u> play in promoting or suppressing the T cell-mediated destruction of pancreatic beta cells in NOD mice (Markle et al. 2013). <u>the possibility that substrain genetic differences may also account for marked variations in diabetes incidences among different colonies of NOD mice has remained an open question</u>. In this regard, NOD males show the greatest variation in diabetes penetrance when compared across colonies, with the microbiome recently demonstrated as a major contributory factor {(Markle et al. 2013).

The advantages and pitfalls of genetic analysis of closely related strains have been recently demonstrated in the comparison of C57BL/6J vs. C57BL/6N (Kumar et al. 2013) and BALB/cJ vs. BALB/cByJ (Sittig et al. 2014). The limited number of polymorphisms between substrains enables their manual curation and increases a chance for identification of single coding polymorphism responsible for the variation of the phenotype. An additional advantage to studying genomic comparisons within NOD substrains is the availability of BAC libraries for two of them, NOD/MrkTac and NOD/ShiLtJ (Steward et al. 2010); https://www.sanger.ac.uk/resources/mouse/nod/).

Here, we report the results of a screen for genetic drift among selected NOD substrains utilizing a customized, high density genotyping chip array combined with whole exome sequencing.

## RESEARCH DESIGN AND METHODS

### Substrains

High molecular weight genomic DNA prepared from NOD/ShiJcl kidney was kindly provided by Dr. K. Hamaguchi (Oita University, Japan.). NOD/ShiLtJ and NOD/ShiLtDvs genomic DNA was prepared from spleens by the JAX DNA Resource and from tail snips of NOD/MrkTac (kindly provided by Dr. L. Wicker, Cambridge University, UK) and the NOD/BomTac substrains (kindly provided by Dr. H.-J. Partke, Diabetes Research Institute, Düsseldorf, Germany). It should be noted that the nomenclature for NOD/LtJ and NOD/LtDvs was changed in 2007 by addition of the source colony descriptor “Shi”; although the MrkTac and BomTac substrains share this common origin, their substrain nomenclatures have not yet been changed to reflect this. The Lt substrain has been bred by Dr. E. Leiter at The Jackson Laboratory since 1984 and sent to its distribution arm in 1992. The Dvs substrain was separated from the Lt substrain in 1992 at The Jackson Laboratory where it is maintained by Dr. D. Serreze as a research colony. The progenitors of the MrkTac substrain were derived from mice from Dr. Y. Mullen’s research colony at UCLA via Dr. L. Wicker (Merck Research Laboratories). The BomTac substrain breeding stock was obtained from a research colony maintained by Dr. L. Herberg (Diabetes Research Institute, Düsseldorf) who received the progenitors from Japan in 1984.

### Exome capture and sequencing

One μg of each substrain DNA was fragmented using Covaris E220 (Covaris, Woburn, MA, USA) to a range of sizes centered on 300 bp. The five pre-capture sequencing libraries were prepared using the NEBNext DNA Library Prep Master Mix Set for Illumina (New England BioLabs, Ipswich, MA, USA) including a bead-based selection for inserts with an average size of 300 bp. The resulting precapture libraries were hybridized to the Roche NimbleGen Mouse Exome capture probe set (Roche NimbleGen, Madison, WI, USA) according to the manufacturer’s instructions. Individually indexed library samples from NOD/ShiLtJ, NOD/LtDvs, and NOD/MrkTac were pooled for exome capture with NOD/Bom and NOD/Shi in a separate capture pool. The two final captured libraries were amplified by 18 cycles of PCR using Phusion High Fidelity PCR mix (NEB). The resulting sequencing libraries were quantified by QPCR, combined, and sequenced on an Illumina HiSeq 2500 (Illumina, San Diego, CA, USA).

Raw reads in FASTQ format were aligned to the mouse reference genome (GRCm38, mm10) using bwa 0.5.9 aligner (Li et al. 2009). SNPs, short insertions and deletions (Indels) were called by the UnifiedGenotyper of Genome Analysis Toolkit 3.1-1 (DePristo et al. 2011), validated by samtools 0.1.19 (Li et al. 2009) with heterozygous calls filtered out, manually checked in IGV 2.1 viewer (Broad Institute) and annotated in Ensembl Variant Effect Predictor. The identified polymorphisms were tested for presence in insulin-dependent diabetes (*Idd*) genetic regions as listed in Table 6.2 of (Ridgway et al. 2008). Note that due to the restriction on gap extension in bwa and the use of UnifiedGenotyper, the chance of identifying larger indels (>5bp) is limited in this analysis. Data are deposited at the Sequence Read Archive (SRA), accession SRP045183.

### Mouse Diversity Array

The high density Mouse Diversity Array (MDA) comprises over 623,000 single nucleotide polymorphism (SNP) probes plus an additional 916,000 invariant genomic probes targeted to genetic deletions or duplications (Yang et al. 2009). DNA was prepared from tissue samples as described above, and genotyped using MDA as previously described (Yang et al. 2009). Raw data (CEL files) are available at ftp://ftp.jax.org/petrs/MDA/raw data, see Suppl. Table 1 for CEL identifications, as implemented in genotypeSnps function of R package MouseDivGeno (Didion et al. 2012), with heterozygous calls filtered out. For detection of copy number variation (CNV), we used simple CNV function of the same package. Only long CNVs covering a number of probes can be discovered by this method (Wang et al. 2007)) ((Bengtsson et al. 2008).

### PCR Validation of Chr. 3 deletion

To validate the substrain-limited deletion on proximal Chr.3, two pairs of PCR primers were designed and the products were Sanger sequenced on an ABI 3730 sequencer at JAX core services. The first pair of primers D3Jmp20 5’-TGTGGTGGACATTTGGGATA-3’ (forward), 5’-AGGCACAGGCAGATCATTCT-3’ (reverse) designed on either side of the approximately 110 kb deletion, gave a 362 bp band from NOD/ShiLt and no band from NON/LtJ. The deletion PCR product was Sanger sequenced to confirm the exact breakpoints. The second pair D3Jmp21 5’-ATCACAGGGTGATCACAGCA-3’ (forward) and 5’-TGTGTTCTTTTCACCCACCA-3’ amplify a product from within the deletion, and gave no product from NOD/ShiLt (region deleted) and a 469 bp band from NON/LtJ.

## RESULTS

### Substrain Genome Comparisons by Exome Sequencing and MDA Analysis

Table 1 shows a comparison of the numbers of SNPs and short indels distinguishing the exomes of the NOD/ShiLt, NOD/ShiLtDvs, NOD/MrkTac, NOD/BomTac and NOD/ShiJcl substrains. The chromosomal locations are provided in Suppl. Table 2 for SNPs and Suppl. Table 3 for short indels. A summary of all polymorphisms and their consequences are shown in Fig. 1. Substrain relationships based upon these polymorphisms were used to estimate a phylogeny tree under generalized time reversible model (Felsenstein 2004) (Fig. 2). This analysis divides the substrains into two clusters; the NOD/ShiLtJ and NOD/ShiLtDvs that were separated most recently and the rest, NOD/ShiJcl, NOD/MrkTac, and NOD/BomTac. Test of the molecular-clock hypothesis, that length of edges is linearly dependent on time of separation, was rejected by a likelihood ratio test (p = 0.007). This could reflect differences in husbandry practices (colony size) resulting in different rates of fixation.

**Table 1:** Number of all SNPs, all indels and coding SNPs + indels distinguishing each pair of substrains are compared to an approximate number of years since separation (exome sequencing only).

|  | All SNP |  |  |  |  | All Indels |  |  |  |  |
| --- | --- | --- | --- | --- | --- | --- | --- | --- | --- | --- |
|  | BomTac | MrkTac | ShiJcl | ShiLtDvs | ShiLt | BomTac | MrkTac | ShiJcl | ShiLtDvs | ShiLt |
| **BomTac** | 0 | 79 | 80 | 86 | 85 | 0 | 22 | 29 | 22 | 26 |
| **MrkTac** | 79 | 0 | 81 | 74 | 73 | 22 | 0 | 25 | 26 | 30 |
| **ShiJcl** | 80 | 81 | 0 | 89 | 88 | 29 | 25 | 0 | 31 | 27 |
| **ShiLtDvs** | 86 | 74 | 89 | 0 | 37 | 22 | 26 | 31 | 0 | 18 |
| **ShiLt** | 85 | 73 | 88 | 37 | 0 | 26 | 30 | 27 | 18 | 0 |
|  | Coding SNP + indels |  |  |  |  | Years since separation (approx.) |  |  |  |  |
|  | BomTac | MrkTac | ShiJcl | ShiLtDvs | ShiLt | BomTac | MrkTac | ShiJcl | ShiLtDvs | ShiLt |
| **BomTac** | 0 | 33 | 33 | 25 | 33 | 0 | 28 | 27 | 27 | 27 |
| **MrkTac** | 33 | 0 | 30 | 22 | 30 | 28 | 0 | 28 | 28 | 28 |
| **ShiJcl** | 33 | 30 | 0 | 24 | 32 | 27 | 28 | 0 | 27 | 27 |
| **ShiLtDvs** | 25 | 22 | 24 | 0 | 14 | 27 | 28 | 27 | 0 | 19 |
| **ShiLt** | 33 | 30 | 32 | 14 | 0 | 27 | 28 | 27 | 19 | 0 |

**Figure 1:**
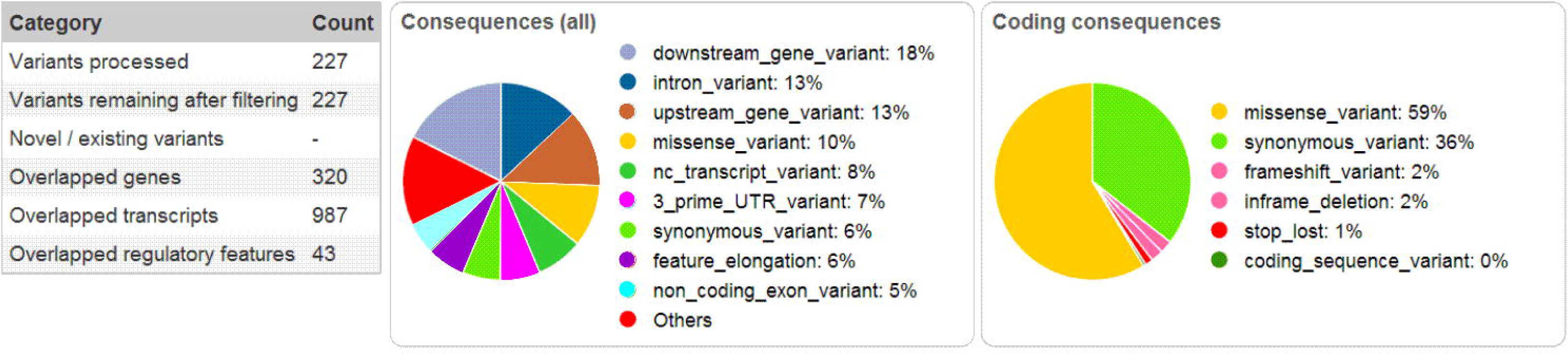
Consequences of polymorphisms identified by exome sequencing categorized by Ensembl Variant Effect Predictor. Totally, 172 SNPs + 55 indels = 227 variants are categorized and percentages of potential consequences is given (A) for all consequences (B) for consequences in coding region. See Ensembl Variant documentation for explanation of consequence categories (http://www.ensembl.org/info/genome/variation/predicted_data.html#consequences)

**Figure 2:**
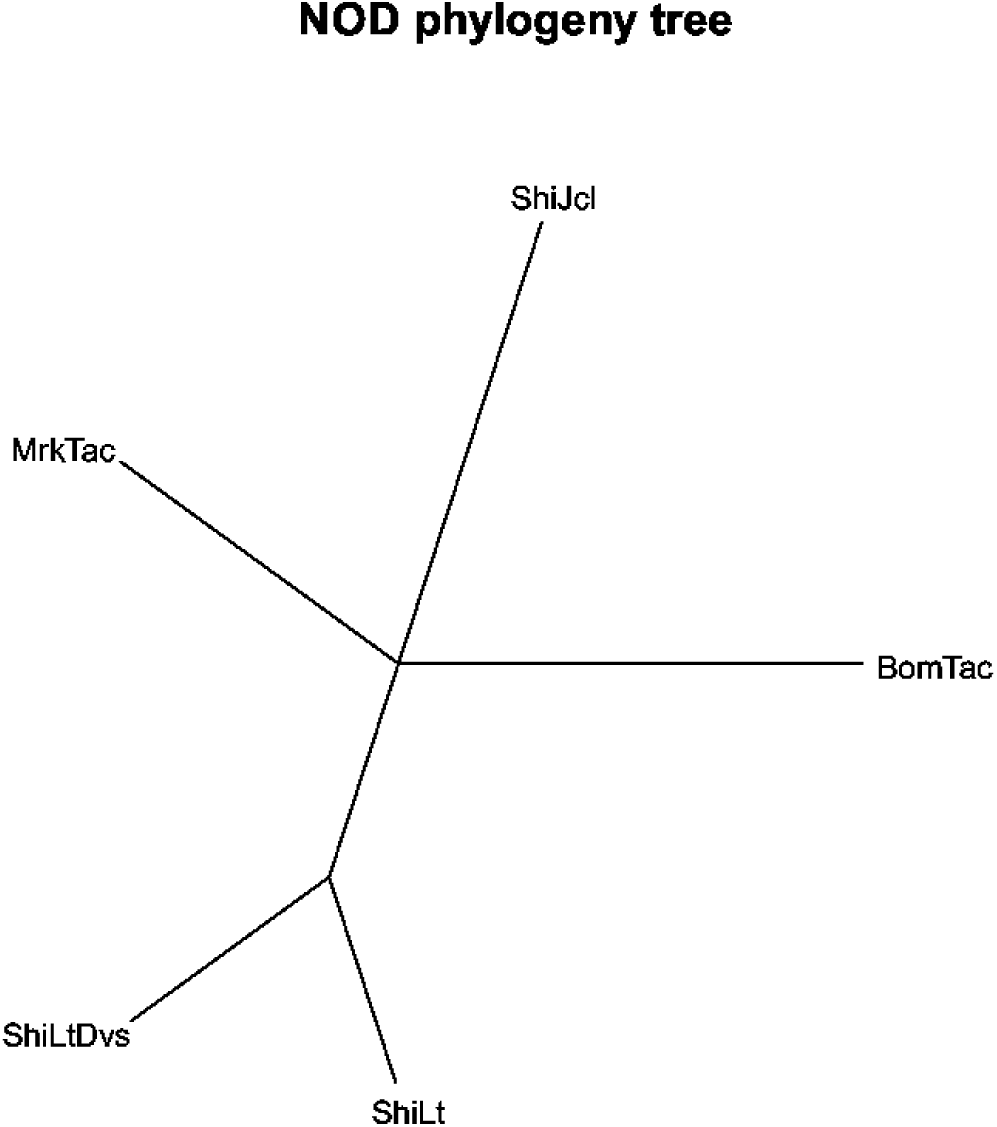
Phylogeny tree of NOD substrains. The length of edges in the tree corresponds to number of SNPs between substrains.

Not surprisingly given their more recent separation, the fewest polymorphisms distinguish LtDvs and LtJ. However, in the relatively short period since these two substrains separated, a deletion occurred in the Dvs *Icam2* (intracellular adhesion molecule2) gene on Chromosome (Chr) 11 that now distinguishes this substrain from both the source LtJ stock and the other three substrains. The loss in intensity of MDA probes only in the Dvs *Icam2* region (Fig. 3) was confirmed by exome sequencing as a deletion that included all exons except the last one (Suppl. Fig. 1). Flow cytometric analysis further confirmed the complete absence of ICAM-2 protein on Dvs substrain leukocytes. Although the *Icam1* gene is essential for diabetes development in NOD (Martin et al. 2001) the presence of this deletion shows that the *Icam2* gene in the Dvs substrain is dispensable for diabetogenesis. Flow cytometric analyses of congenic stocks built using NOD/ShiLtDvs as the recipient strain indicated those initiated before and after approximately the year 2000 are respectively ICAM-2 intact and deficient. This indicates the *Icam2* deletion presently characterizing the NOD/ShiLtDvs substrain occurred around the year 2000.

**Figure 3:**
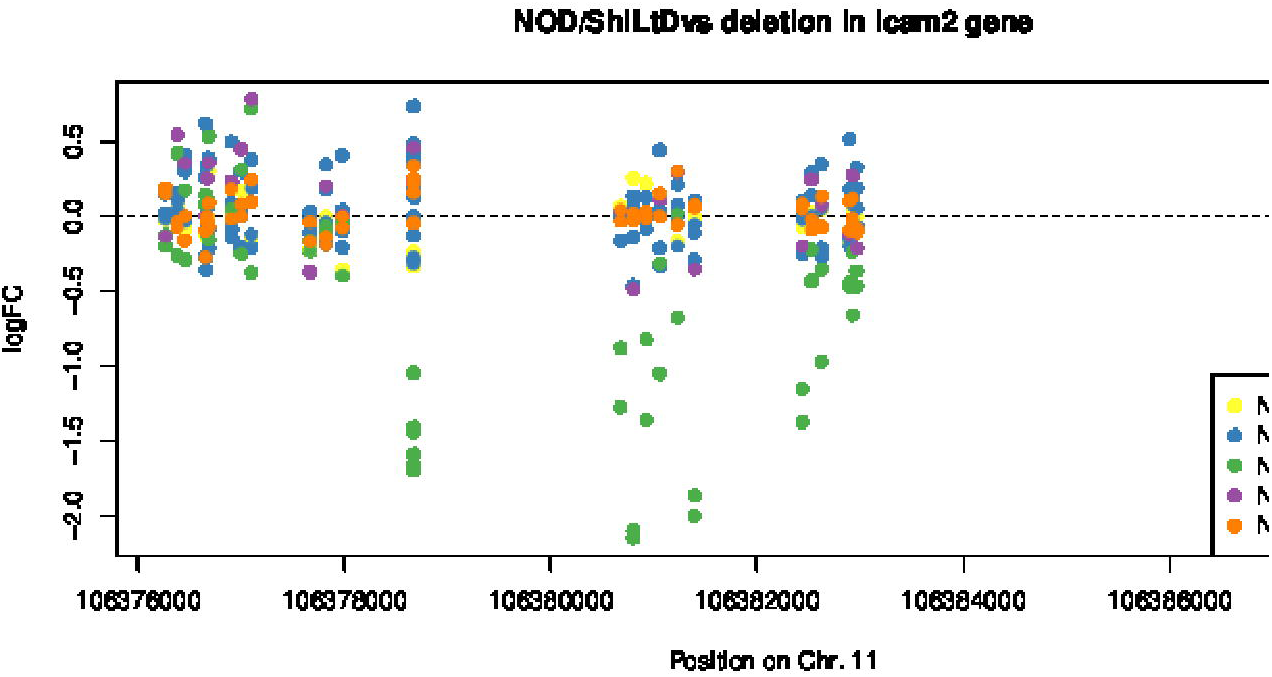
Scaled intensities of MDA probes along the *Icam2* region. Negative NOD/ShiLtDvs values suggest a deletion. See also Suppl. Fig. 1.

Compared to exome sequencing, MDA enables discovery of specific SNPs in noncoding regions and copy-number variations. In total, 48 SNPs and seven CNVs were detected, listed in Suppl. Tables 4 and 5. Among these, a 110,275 bp deletion on proximal Chr. 3 was unique to the LtJ and Dvs substrains. This deletion in a gene-poor area spanned 13 SNPs between 24,272,052 and 24,383,100 bases and was validated by Sanger sequencing across the deletion in NOD/ShiLt (see Methods). The only protein coding gene at this interval is *Naaladl2* (N-acetylated alpha-linked acidic dipeptidase-like 2) whose expression in mouse is primarily limited to the eye. No overt eye phenotype has been observed to differentiate these substrains. SNPs covering this deletion provide an excellent means for distinguishing the closely-related LtJ and Dvs substrains from the other three more genetically divergent substrains.

Supplementary Tables 2 and 3 provide substrain distribution patterns from the whole exome sequencing data for SNPs and indels respectively. We identified 64 coding SNPs and 2 indels. Of those SNPs that effect a missense change in protein coding exons, many are associated with neuronal functions (for example, the CNS-restricted protein tyrosine phosphatase, receptor type 2 mutation (*Ptprt*, Chr.2) limited to the LtJ substrain). Some missense mutations are unique for a given substrain; e.g., missense mutations in *Tmem163* and *Fam129b* (Chr.1), *Tet2* (Chr.3), *Cpz* and *Nsg1* (Chr.5), *Prdm10* (Chr.9), *Arsg* (Chr.11), *Pcnx* and *Akr1c12* (Chr.12), *Liph* (Chr.16), and *Exo6* (Chr.18) are limited to BomTac. The missense mutations are differentially distributed across the five substrains, so that it is not possible to predict whether any one of them alone affects a critical immunophenotype affecting diabetes penetrance. This was also the case for the substrain distribution of insertions and deletions summarized in Supplementary Table 3. Whereas the locations and size of the indels were most closely matched in the closely-related LtJ and Dvs substrains (each with 9 deletions and 11 insertions), they were not all concordant across the genome. Comparison of the indels in the other 3 substrains indicated the independence of these genetic changes.

### PCR jumping

Sequencing reads from a given substrain at a specific SNP are expected to show close to 100% base call identity, consistent with the inbred nature of the strains. However, we also analyzed the frequency of rare allele calls, as a measure of sequence error rates or possible contamination between samples.

From the 172 SNP loci we selected 22 loci with high coverage [all five strains had at least 100 sequencing reads covering that base position (Suppl. Table 6)]. There were no consistent sources of rare allele calls that would implicate sample or barcode contamination. Sequencing error rate was estimated as 0-0.33% (mean 0.05). However, for a given substrain, the presence of a minor variant (present as a major allele in a different substrain in the same exome capture pool) was observed at a significantly higher frequency, 0.7-3.8% (mean 1.6%). Every locus with more than 100 reads in all 5 strains showed this higher rate of rare allele calls within pools. This observation is consistent with previous reports of “Jumping PCR” (Ramos et al. 2012; Kircher et al. 2012). This phenomenon occurs during post-capture PCR library amplification, wherein a nascent DNA strand switches templates from one allele to another. This “Jumping PCR” generates apparent rare allele calls that could be problematic in interpretation of sequence data from outbred animals or human populations, but for sequencing studies using inbred mice, this type of low frequency artifact is readily distinguishable from the expected homozygous allele calls.

## DISCUSSION

In the 34 years since the original report of the NOD/Shi strain in Japan, the genomes of the derivative colonies are clearly exhibiting genetic drift. From the results shown in Table. 1, we can estimate that number of coding polymorphisms for a year of separation among NOD substrains is 0.74-1.22 (pair-wise comparisons among 5 substrains), or 0.19-0.31 for a generation (under assumption of 4 generations per year). Our results are similar to a recent paper (Simon et al. 2013) comparing C57BL/6J and C57BL/6N mouse strains separated around 220 generations ago that identified 34 SNPs and 2 indels in the coding region.

One of the major phenotypic differences distinguishing various colonies of NOD mice worldwide has been the penetrance of diabetes, particularly in males. With the exception of the low diabetes incidence NOD/Wehi substrain where a single recessive mutation may explain this now extinct substrain’s diabetes resistance (Baxter et al. 1993), the genetic basis for differential diabetes penetrance in the currently distributed NOD substrains, if any, is unknown. Although our study shows that each substrain carries unique indels or missense mutations that distinguish them from one another, no single one of these mutations has a known effect on diabetes penetrance or an immunophenotype related to it. The Chr. 3 deletion in a gene poor region in LtJ and Dvs is at least 1 Mb proximal to the *Idd3/Il2* locus known to be a major determinant of diabetes susceptibility in NOD mice (Yamanouchi et al. 2007). As noted in Methods, in order to avoid any ambiguity resulting from false calls, this analysis was exclusively focused on homozygous calls. Hence, this analysis has uncovered only a subset of all the genomic differences that must distinguish these 5 substrains as their genomes continue to drift over time (for example, recent heterozygous mutations not yet fixed to homozygosity).

In conclusion, we have documented genetic drift among NOD substrains that allows for distinguishing them genetically when necessary. The extent to which the polymorphisms identified potentially contribute to phenotypic differences <u>among substrains</u> remains unclear. This study only focused on variants fixed to homozygosity; additional heterozygous mutations likely are continuing to be fixed to homozygosity with successive generations of inbreeding. That such mutations may affect phenotype is clear. For example, a cohort of NOD/Jos mice received by other investigators in 1988 had diverged into high and low diabetes incidence sublines by 1993 (Takayama et al. 1993). Cryopreservation efforts to keep genetic drift to a minimum are clearly useful in maintaining a consistent phenotype (Taft et al. 2006). <u>Finally, the existence of genetic divergence among NOD substrains emphasizes the importance of using the appropriate genetic nomenclature that permits identification of NOD colonies that have been separated from a source colony for 10 generations of inbreeding or more.</u>

## ACKNOWLEDGEMENTS

The authors thank Drs. Kazuyuki Hamaguchi, Linda Wicker, and Hans-Jochim Partke for kindly supplying substrain DNA as noted in the Methods. The work was also supported by NIH grants P50GM076468 (GAC), DK095735, DK046266, UC4-DK097610, grant 2014PG-T1D048 from the Helmsley Charitable Trust, and grants from Juvenile Diabetes Research Foundation and The American Diabetes Association (DVS). Shared Services of The Jackson Laboratory used in this publication was partially supported by the National Cancer Institute grant P30CA034196. The content is solely the responsibility of the authors and does not necessarily represent the official views of the NIH.

## Figures and Tables

**Supplemental Figure 1:** IGV Browser screen with the coverage of exome sequencing in the *Icam2* region. All but the last *Icam2* exon (ENSMUSE00000374954) are missing in the NOD/ShiLtDvs genome.

**Supplemental Table 1:** MDA and exome sequencing file identifiers and quality measures: number of reads, target fold coverage and percentage of target covered with >= 10 reads (exome sequencing); percentage of MDA probesets’ calls (2 B6 alleles / 2 non-B6 alleles / heterozygous / drop in intensity / no call)

**Supplemental Table 2:** List of SNPs among substrains identified by exome sequencing annotated by Ensembl Variant Effect Predictor. Coding SNPs are highlighted in bold. If the SNP is covered by more than one gene or transcript, all possible consequences are listed (separated by “//”).

**Supplemental Table 3:** List of short indels identified by exome sequencing annotated by Ensembl Variant Effect Predictor. Coding indels are highlighted in bold. If the SNP is covered by more than one gene or transcript, all possible consequences are listed (separated by “//”).

**Supplemental Table 4:** List of SNPs and VINOs from Mouse Diversity Array analysis annotated by Ensembl Variant Effect Predictor. If the SNP is covered by more than one gene or transcript, all possible consequences are listed (separated by “//”)

**Supplemental Table 5:** List of (long) deletions and extra copies identified by Mouse Diversity Array copy number variation analysis.

**Supplemental Table 6:** Rare allele frequency in 5 NOD substrains. Only loci with >100 reads for each strain are included. Grey cells indicate variants that were not represented as a predominant allele in any of the 5 strains (included in the % sequencing error calculation), pink cells are variants that matched the predominant allele in one or more strains in the opposite PCR pool (included in the inter-pool error calculation), and yellow cells are variants matched the major allele of other substrains in the same PCR pool (included in the itra-pool jumps calculation). Note that some allele data fall into more than a single class. Red text indicates class (1) and (2) variants at >1 rare allele read count.

